# Anxiety-related attentional characteristics and their relation to freezing of gait in people with Parkinson’s – cross-validation of the Adapted Gait Specific Attentional Profile (G-SAP-PD)

**DOI:** 10.1101/2024.03.14.585018

**Authors:** Uri Rosenblum, Adam J. Cocks, Meriel Norris, Elmar Kal, William R. Young

**Author notes:** Corresponding Authors: Uri Rosenblum, PhD PT, Department of Physical Therapy, College of Health, Medicine and Life Sciences Brunel University London, Uxbridge, UB8 3PH, United Kingdome. Joint senior authorship (Elmar Kal, William R. Young).

## Abstract

**BACKGROUND:** Anxiety often exacerbates freezing of gait (FOG) in people with Parkinson’s (PwP). Research shows that anxiety-related attentional processes and associated processing inefficiencies, such as conscious movement processing (CMP) and ruminations, can substantially impact movement control. However, the impact of these attentional characteristics on FOG remains largely unexplored.

**OBJECTIVES:** To (i) validate an adapted 10-item (1-5 Likert scale) Gait-Specific Attentional Profile in PwP (G-SAP-PD), and (ii) assess if G-SAP-PD-subscales (Physiological Arousal, CMP, Rumination, and Processing Inefficiencies) are associated with self-reported FOG frequency.

**METHODS:** We recruited 440 PwP (M_age_=65.5±8.7; 5.8±5.0 years since diagnosis) across the UK. Participants completed the G-SAP-PD, and questions on demographics, medical background, and FOG frequency (scale of 0: “never freeze” to 4: “every day”). We assessed G-SAP-PD’s internal consistency (alpha), structural validity (confirmatory factor analysis), and subscale scores associations with FOG frequency (ordinal regression).

**RESULTS:** The G-SAP-PD’s showed high internal consistency (α>0.61) and acceptable/good model fit (comparative fit index=0.976). Physiological Arousal and CMP subscale scores were less strongly correlated for PwP with FOG (PwP+FOG, r=.52, p=0.001) compared to (PwP-FOG, r=.79; p=0.001). Higher Rumination (OR: 1.323, 95%CI: [1.214-1.440]) and Physiological Arousal (OR: 1.195, 95%CI: [1.037-1.377]) were significantly associated with higher FOG frequency, when controlling for age, time since diagnosis and balance/gait problems.

**CONCLUSIONS:** The G-SAP-PD is a reliable and convenient tool to measure and identifying potentially maladaptive anxiety-related attentional processes that might impact FOG. Our data suggests a relative inability of PwP+FOG to engage in compensatory goal-directed attentional focus. Further study is warranted.

**Plain Language Summary:** Anxiety can worsen freezing of gait in people with Parkinson’s. It often leads to worrisome thoughts, and influences how people pay attention to their walking. We think that these changes in attention can substantially influence peoples’ movement – for better or worse. However, there is a lack of research on this topic, and reliable assessment tools are missing.

Therefore, we tested if we could assess changes in the thoughts and attention of people with Parkinson’s, using a questionnaire (Gait-Specific Attentional Profile (G-SAP-PD)), previously used in older adults without Parkinson’s. This questionnaire aims to measure people’s perception of their physiological arousal (how anxious they feel), conscious movement (attention they direct to walking), rumination (worrisome thoughts), and thinking efficiency (the ability to focus on different tasks). We also investigated if people who experience freezing show different attentional characteristics compared to people who do not experience freezing. Four-hundred and forty people with Parkinson’s filled the G-SAP-PD questionnaire. We confirmed the questionnaire’s reliability, and found that people who indicated to have more worrisome thoughts and greater physiological arousal also experienced freezing more often. Our findings suggest that people with Parkinson’s who experience freezing were less able to consciously direct attention to the task at hand (taking a step) when experiencing high physiological arousal. The G-SAP-PD represents a short and convenient tool for identifying potentially negative attentional and thinking processes that may increase freezing frequency. With further research it could be used as a predictive tool and provide possible novel treatments to reduce freezing frequency.

## Introduction

Freezing of gait (FOG) is a “brief episodic absence or marked reduction of forward progression of the feet despite the intention to walk”[1]; a debilitating symptom of Parkinson’s, with prevalence reported as high as 80%[2]. FOG is associated with increased prevalence of falls[3] and reduced quality of life of People with Parkinson’s (PwP)[4,5]. FOG can manifest across different so-called phenotypes, such as trembling in place, shuffling forward, and akinesia. While there currently is no broad consensus on the specific pathogenesis of FOG, common triggers have been identified, such as doorways/narrow spaces, turning and multi-tasking[6,7]. One specific factor that is often implicated in exacerbating FOG frequency and duration is anxiety[8]. It is suggested that increased anxiety (i.e., physiological arousal) overwhelms pre-existing defective limbic circuitry and noradrenergic pathways, leading to neural and cognitive inefficiencies that worsen freezing[8,9]. Based on the Attention Control Theory (ACT)[10], this paper aims to study the role of specific anxiety-related attentional characteristics in FOG, as this may ultimately help inform intervention development.

ACT[10] describes how anxiety may lead to preferential engagement of the ventral stream of information processing and associated <u>stimulus-driven</u> attentional system at the expense of a decreased influence of the <u>goal-directed</u> (dorsal) attentional system[10]. According to these predictions, PwP with high arousal would allocate attentional resources to processing threat-related stimuli (e.g., worrisome thoughts related to freezing, or threatening task-irrelevant distractors, like doorways) rather than to goal-directed attentional processes (e.g., focusing on the intended step), resulting in more frequent and severe freezing. However, based on ACT, and its predecessor Processing Efficiency Theory[11], it is also predicted that such negative effects of anxiety could be offset if PwP manage to maintain their attentional focus toward the intended movement goal, through increased mental effort and/or inhibition of distraction by threat-related stimuli. These theories are well supported by empirical evidence in performance contexts, such as sport or surgery[12–14] as well as functional gait, especially in older adults at risk of falling[15]. However, relatively little is known about the role of these proposed anxiety-related attentional characteristics in the context of FOG. There is evidence that worrisome thoughts and rumination (i.e., self-preoccupation with concerns over failure and expectations of negative consequences[16] are triggered by stressful situations and are prevalent in individuals high in trait anxiety[17] such as older adults fearful about falling. While such worries may be acted on to make adaptive changes to behaviour (e.g., walking with an assistive device), processing such thoughts while walking will be cognitively demanding, and will bias an individual’s attention toward potential threats to balance[18,19] (or in PwP: toward potential triggers for FOG). This may distract attention away from the movement task at hand, and thus exacerbate FOG. These assertions are particularly relevant in the context of Parkinson’s where increased conscious monitoring of ongoing movements is required to compensate for loss of movement automaticity[20].

The negative effects of anxiety-induced rumination and worry may therefore be minimised by investing more conscious attention into controlling and monitoring ongoing movement. Nonnekes and colleagues[21] identified 59 unique strategies for improving mobility in PwP, “where an overarching working mechanism involved in all was allocation of attention to gait, the introduction of goal directedness, and the use of motor programs that are less automatized than those used for normal walking”[21]. Further, cueing and movement strategy (e.g., weight-shifting) interventions for FOG are thought to be effective, at least in part, because they compensate for deficient automaticity by engaging cortical networks involved in goal-directed attention[22,23]. However, conscious goal-directed strategies are effortful and cognitively demanding, and will become more difficult to employ as anxiety and overall task demands increase[10]. This may be especially true for PwP, who often already demonstrate both strong conscious control of movement[24], as well as deficits in executive functions, such as inhibition[25–29] and shifting of attention[30–36]. This will likely compromise their ability to block out worrisome thoughts and shift attention back toward the movement task at hand.

The Gait-Specific Attentional Profile (G-SAP), a short self-report instrument, has recently been developed for older adults with balance impairments. The instrument allows the measurement of the degree to which individuals experience heightened somatic anxiety (Physiological Arousal subscale), conscious attention to movement (Conscious Movement Processing, CMP subscale), worrisome thoughts (Rumination subscale), and processing inefficiencies (PI subscale) when walking in daily life[18]. The G-SAP could provide insight into the role of these different constructs in the context of FOG in PwP. Indeed, a recent study by Cockx et al.[37] used the G-SAP to explore potential relationships between the above constructs and the propensity to experience FOG when navigating doorways. They found that people with longer disease duration and who show freezing in response to doorways have significantly higher scores for all G-SAP subscales compared to those who do not freeze in response to doorways. While this report does not identify freezing pathology as an independent factor associated with higher G-SAP scores, the results clearly emphasise the potential of utilising self-reported attentional processes to deepen our understanding of the relationship between anxiety and FOG in different contexts. Further, a limitation of Cockx et al.[37] is that no comprehensive cross-validation has yet been conducted to demonstrate the reliability and validity of the GSAP for PwP with (PwP+FOG) and without FOG (PwP-FOG). Lastly, the original G-SAP questionnaire consists of 11 questions across the same four domains. However, item A2, from the ‘Physiological Arousal’ sub-scale, does not fully capture/align with the subscale of Physiological Arousal. That is, item A2 required respondents to indicate to what extent they are concerned about other people’s thoughts about them. While this item was allocated to the more broadly named ‘anxiety’ sub-scale in the original G-SAP validation (in a heterogeneous group of older adults), there is a concern that this concept does not fit with the more specific construct that better-reflects physiological arousal in the context of PwP.

Therefore, this study aimed to 1) validate an adapted version of the G-SAP scale for use in PwP+FOG and PwP-FOG (G-SAP-PD), where item A2 in the original G-SAP was removed from the ‘Physiological Arousal’ sub-scale. This has the benefit of creating a clearer contrast between Physiological Arousal (A1,A10) and Rumination (A3,A4,A6) ; and 2) determine if self-reported FOG frequency in daily life is independently associated with different G-SAP-PD sub-scales. In line with the literature above[17,37], we hypothesised that more frequent FOG would be associated with higher scores on all subscales, but would show strongest unique associations with Rumination subscale scores.

## Methods

### Participants

Four hundred and forty PwP were recruited through advertisements across the Parkinson’s UK network (circulated through email across the UK) as well as in-person invitations at local Parkinson’s support groups in West London. The advertisements included a link to an online survey. No incentives were provided for participation.

Participants were eligible for inclusion if they had a diagnosis of Parkinson’s and had sufficient command of the English language to understand and complete the survey. Participant characteristics are presented in Table 1.

**Table 1.** Characteristics of patients with (‘PwP+FOG) and without (‘PwP-FOG’) freezing of gait.

|  | **PwP-FOG (N=236)** | **PwP+FOG (N=199)** |  |  |
| --- | --- | --- | --- | --- |
|  | *"Never"*<br>(N=236) | *"Hardly ever"*<br>(N=109) | *"Most weeks"*<br>(N=39) | *"Every day"*<br>(N=51) |
| **Age in years** | 65.9±8.2<br>[43-81]<sup>c</sup> | 63.3±8.0<br>[40-81]<sup>e</sup> | 64.3± 10.9<br>[41-85]<sup>b</sup> | 68.7±9.0<br>[46-86]<sup>a</sup> |
| **Any other diagnosis/condition<sup>#</sup>** |  |  |  |  |
| *None* | 186 (79%) | 75 (69%) | 30 (77%) | 35 (69%) |
| *Arthritis* | 16 (7%) | 10 (9%) | 3 (8%) | 6 (12%) |
| *Neurological* | 7 (3%) | 6(5%) | 2 (5%) | 1 (2%) |
| *Back pain/problem* | 9 (4%) | 6 (5%) | 1 (3%) | 2 (4%) |
| *Other orthopaedic* | 12 (5%) | 5 (5%) | 1 (3%) | 2 (4%) |
| *Cardiovascular* | 2 (1%) | 2 (2%) | 0 (0%) | 4 (8%) |
| *Other (e.g. diabetes, colitis, tendon injury)* | 2 (1%) | 5 (10%) | 1 (3%) | 0 (0%) |
| **Years since diagnosis** | 4.3±3.4***<br>[0-19]<sup>a</sup> | 5.8±4.6<br>[0-23]<sup>d</sup> | 8.2±5.8<br>[0-25]<sup>a</sup> | 10.5±7.1<br>[0-27] |
| **Tremor** | 2 (1) | 2 (1) | 2 (2) | 2 (2) |
| *Not at all (1) - Very much so (5)* | [1-5]<sup>a</sup> | [1-5] | [1-4]<sup>a</sup> | [1-5] |
| **Problems with balance and gait** | 2 (1)***<br>[1-5] | 2 (2)<br>[1-5] | 3 (2)<br>[1-5]<sup>b</sup> | 4 (2)<br>[1-5] |
| *Not at all (1) - Very much so (5)* |  |  |  |  |
| **G-SAP-PD Sub-scale Scores** |  |  |  |  |
| Physiological Arousal | 5.0±1.9***<br>[2-10] | 6.4±2.0<br>[2-10] | 6.9±1.9<br>[3-10] | 7.4±1.8<br>[4-10] |
| CMP | 9.5±3.0***<br>[3-15] | 10.7±2.6<br>[4-15] | 11.1±2.5<br>[5-15] | 11.9±2.2<br>[7-15] |
| Rumination | 5.6±2.5***<br>[3-13] | 7.8±2.8<br>[3-15] | 8.4±2.7<br>[3-13] | 10.8±2.8<br>[3-15] |
| PI | 3.8±1.7***<br>[2-9] | 5.0±1.7<br>[2-9] | 5.2±2.0<br>[2-9] | 5.8±2.1<br>[2-10] |

**NB:** Continuous variables are expressed as mean ± standard deviation [range], while categorical variables are expressed as median (interquartile range) [range]; #Note that some patients reported more than one additional condition, and that percentages will therefore add up to more than 100%; \*\*\* significant difference between PwP+FOG and PwP-FOG p<0.001;
<sup>a</sup> – one missing value; <sup>b</sup> – two missing values; <sup>c</sup> – four missing values; <sup>d</sup> – five missing values; <sup>e</sup> – eight missing values
Abbreviations: CMP = Conscious Movement Processing; PI = Processing Inefficiency

Institutional ethical approval was obtained from the College of Health, Medicine and Life Sciences Research Ethics Committee of Brunel University London (REF: 6473-MHR-May/2017-7263-2). All participants provided online written informed consent prior to participation.

### Procedure

The online survey was hosted online using JISC Online Surveys (Bristol, UK). First, participants completed the online informed consent form, prior to completing several questions on their background. These included age in years, co-morbid diagnoses (e.g., orthopaedic, neurological conditions), years since Parkinson’s diagnosis, information on other Parkinson’s symptoms such as tremor and balance problems; rated on 1 (“not at all”) to 5 (“very much so”) Likert scale), and self-reported FOG frequency (“never”, “hardly ever”, “most weeks”, “every day”). For the purpose of later analysis (see below), participants who reported to “never” freeze were categorized as PwP-FOG, while all others reporting higher scores were classified as PwP+FOG.

### G-SAP-PD questionnaire

Next participants completed an adapted G-SAP-PD questionnaire (see Methods section in *Supplementary materials)*. This G-SAP-PD consists of 10 questions across 4 different domains: Physiological Arousal (2 items), CMP (3 items), Rumination (3 items), PI (2 items). For each item, respondents indicate their level of agreement on a scale from 1 (“Not at all”) – 5 (“Very much so”). For each subscale, scores are then summed to create an overall subscale score.

### Data analysis & statistics

All data were analysed with R version 4.2.2., unless stated otherwise. Alpha was set at 0.05.

#### Comparison of characteristics between PwP+FOG and PwP-FOG

Patient characteristics were presented using appropriate measures of central tendency and dispersion, and compared between PwP+FOG and PwP-FOG using parametric and/or non-parametric tests as appropriate. Cohen’s d was used as measure of effect size [38] (‘cohensD’ function, ‘lsr’ package in R).

#### Validity and reliability of Gait-Specific Attentional Profile

We conducted confirmatory factor analysis (maximum likelihood estimation, CFA) to assess the G-SAP-PD’s structural validity (AMOS, version 26; IBM, Chicago, IL). Specifically, we aimed to determine whether the data would fit the four-factor structure reported in the initial G-SAP validation study in healthy older adults[18]. Pairs of error terms for items loading on same subscale/factor were allowed to co-vary if this improved model fit.

We present the overall model including standardized item-factor loadings, along with the following model fit tests: Chi-square statistics, both absolute (χ^2^; non-significant χ^2^indicates acceptable fit) and divided by degrees of freedom (χ^2^/df; values<3 indicate acceptable fit); goodness-of-fit and comparative fit indices (GFI and CFI; values>0.90 indicate acceptable fit, values>0.95 indicate good fit); standardized root mean squared residual (SRMR; values<0.008 indicate good fit); and root mean square error of approximation (RMSEA; values<0.05 indicate good fit, values<0.08 indicate acceptable fit[39–41].

Next, we performed measurement invariance tests to determine if the G-SAP-PD’s factor structure is similar for PwP+FOG and PwP-FOG. This consisted of three different steps. First, model fit was assessed when item-factor loadings were free to differ between PwP+FOG and PwP-FOG (configural invariance), subsequently with item-factor loadings fixed across these two groups (metric invariance), and finally with both the item-factor loadings and intercepts fixed (scalar invariance). Factor structure is considered similar if model fit remains similar – i.e., non-significant Δ χ^2^ values, ΔCFI & ΔRMSEA<0.010, ΔSRMR<0.015[42].

Finally, internal consistency was determined for each separate subscale/factor of the G-SAP-PD using ‘alpha’ function in the ‘psych’ package in R. Standardized according to the correlations Alpha ≥ 0.65 was considered to be sufficient internal consistency[43,44].

#### Relation between Gait-Specific Attentional Profile and freezing of gait

Two-Way ANOVA (‘aov’ function, ‘stats’ package in R) was used to compare G-SAP-PD scores between groups of patients with different freezing frequency, with the different subscale scores serving as dependent variable and the independent variables being the G-SAP-PD subscales, freezing frequency, and their interaction (G-SAP-PD subscale X freezing frequency). For this analysis, G-SAP-PD scores were Z-transformed to allow for comparison between subscales with different score range. Post-hoc Tukey Honest Significant Difference was used to correct for multiple comparisons. Eta squared (ƞ^2;^ ‘etaSquared’ function in the ‘lsr’ package in R) was presented as measure of effect sizes.

Subsequently, we performed ordinal regression analysis to analyse the association between the different G-SAP-PD subscale scores and frequency of freezing (logit link function). Participant characteristics (i.e., age, years since diagnosis and experiencing balance problems) were controlled for as covariates in the model.

We then selected the G-SAP-PD subscale score with the highest (significant) odds ratio in the ordinal regression to further explore diagnostic accuracy. Specifically, we used area under the curve (AUC) analysis (SPSS version 29; IBM, Chicago, IL) to determine cut-off scores based on optimal sensitivity vs specificity trade-off (Youden’s index[45]) for (i) PwP+FOG vs PwP-FOG status and (ii) freezing everyday vs freezing less frequently (combined freezing “hardly ever” and “most days”).

#### Sample size calculation

We aimed for an overall sample of ∼400 participants, which is the recommended sample size for factor analysis involving 3-6 factors (subscales), 3 items per factor, and conservative expected factor loadings of 0.4[46]. This sample size was also anticipated to result in at least 30 participants for each freezing frequency category, as required for the 2-way-ANOVA and the ordinal regression model[47,48].

## Results

### Participants

In total, 440 patients with Parkinson’s disease completed the questionnaire. Five were excluded from the CFA due to missing freezing data, leaving 435 patients. Twenty-seven patients were excluded only from the regression analysis, because of missing data regarding freezing (N=5), years since diagnosis (N=9), and/or age (N=16). Thus, regression analysis was performed on the remaining 413 patients. Detailed patient characteristics can be found in Table 1.

### PwP+FOG vs. PwP-FOG

In all, 236 patients reported that they never experience freezing of gait (PwP-FOG), while 199 patients reported to experience freezing (PwP+FOG). On the whole, PwP+FOG were of similar age as PwP-FOG (64.9±9.3 vs. 65.9±8.2, respectively; t_(377.78)_ = 1.11, Cohen’s d=0.11, p=0.867). However, PwP+FOG had longer time since diagnosis of Parkinson’s disease (7.49±5.91 vs. 4.3±3.4, respectively; t_(295.05)_ = -6.63, Cohen’s d=0.67, p<0.001), more self-reported balance problems (Mdn = 3, IQR=2 vs. Mdn=2, IQR=1, respectively; W = 11841, Cohen’s d=0.95, p<0.001). However, presence of tremor complaints (Mdn = 2, IQR=2, vs. Mdn = 2, IQR=1; W = 23639, Cohen’s d=0.00, p=0.619 and of additional medical conditions (30% vs.21%; χ2_(1)_=2.253, Cohen’s d=0.20, p=0.133) were similar for PwP+FOG and PwP-FOG, respectively.

### Gait-specific attentional profile validation for PwP

#### Structural validity: Confirmatory factor analysis (CFA)

The overall CFA model is presented in Figure 1. Medium to strong correlations were observed between all factors, and especially between the factor of ‘Physiological Arousal’ and ‘Conscious Movement Processing’ (r=0.713), Rumination (r=0.700) and Processing inefficiency (r=0.704). Standardised item-factor loadings were all positive and high (≥0.70). Model fit was acceptable to good (χ^2^ =82.833, *p*<0.001; χ^2^/df=2.856; CFI=0.976; GFI=0.963; RMSEA=0.066[0.049, 0.083]; SRMR=0.035).

**Figure 1.**
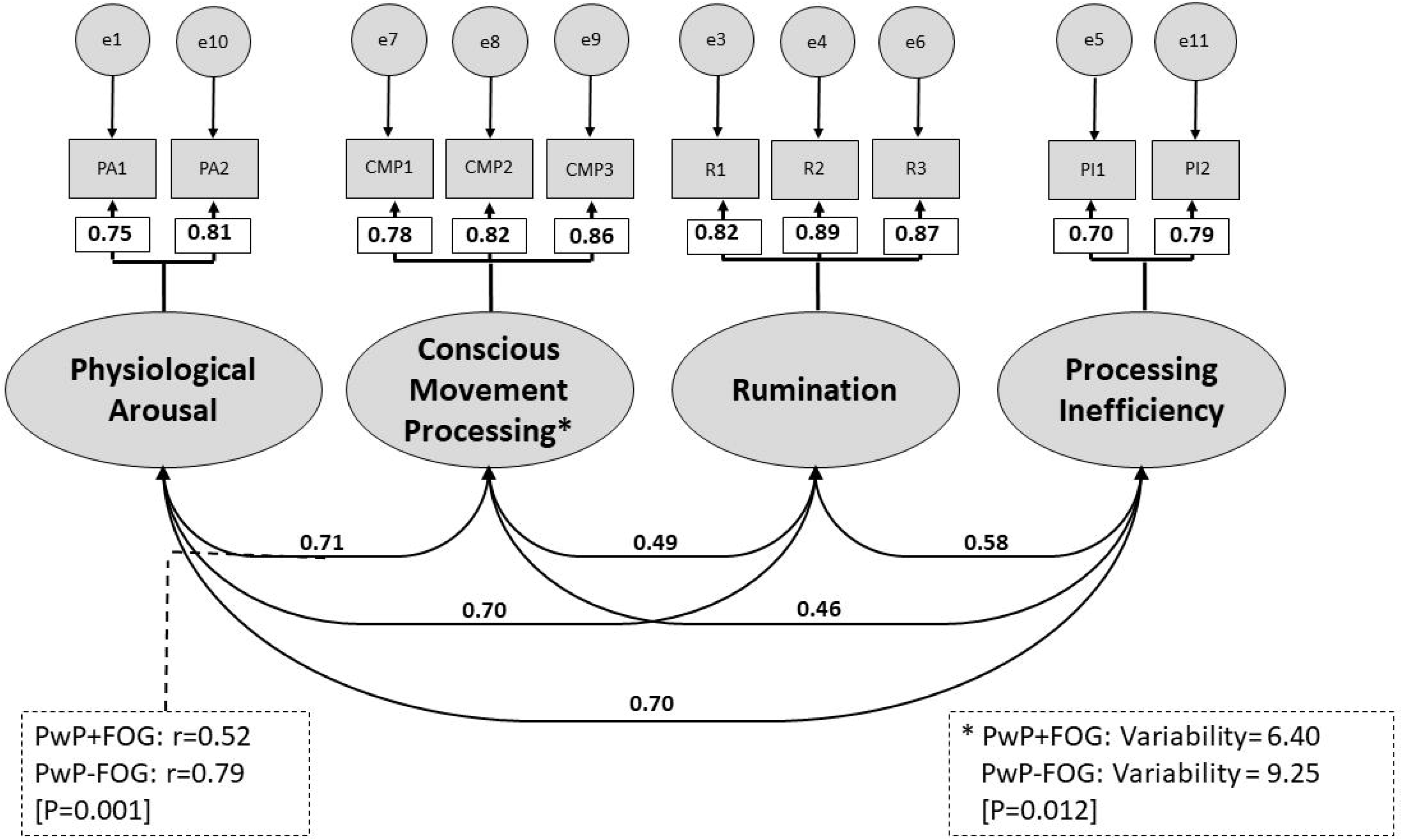
Final overall configural model yielded by the Confirmatory Factor Analysis. Shown are the standardised item-factor loadings and the covariances between the latent factors for all participants combined and specifications of significant differences between people with Parkinson’s with (PwP+FOG) and without freezing of gait (PwP-FOG) in an adjusted partial scalar model. Dotted lines indicate that covariances are significantly different between PwP+FOG and PwP-FOG, with the respective values for both groups. *indicates significant variability differences between PwP+FOG and PwP-FOG, with the respective values for both groups. Abbreviated item numbers refer to the respective items for each latent factor on the Adapted Gait Specific Attentional Profile (G-SAP-PD, see Appendix). NB: PA = Physiological Arousal; CMP = Conscious movement processing; PI = Processing inefficiency; R = Rumination; e = residual error. See Table S1 in *Supplementary materials* for details on final model selection.

The model demonstrated configural and metric measurement invariance (Table S1, in *Supplementary materials*). Scalar measurement invariance was borderline satisfactory. Further analysis using backward releasing of constraints revealed that this was primarily due to a significant between-group difference in covariance between the subscales ‘Physiological Arousal’ and ‘CMP’ (*r*_PwP+FOG_=0.52, *r*_PwP-FOG_=0.79; *Z*=-2.934, *p*=0.003), and to a lesser extent to reduced variability in ‘CMP’ subscale scores among PwP+FOG (6.40) compared to PwP-FOG (9.25; *Z*=-2.254, p=0.012). Unconstrained values for the ‘Physiological Arousal’ and ‘CMP’ covariance, and ‘CMP’ variance led to acceptable partial scalar invariance (Table S1).

Overall, the CFA therefore confirmed the hypothesized four-factor structure of the G-SAP-PD, and demonstrated that the scale was suitable to compare scores between PwP+FOG and PwP-FOG.

#### Internal consistency

Internal consistency of the G-SAP-PD was confirmed. For the sample overall, standardized Cronbach’s alpha values were 0.75, 0.86, 0.89 and 0.71 for Physiological Arousal, CMP, Rumination and PI, respectively. For PwP+FOG the respective standardized Cronbach’s alpha values were 0.69, 0.81, 0.86, and 0.66, while for PwP-FOG these were 0.70, 0.88, 0.85, and 0.68.

### G-SAP-PD scores - differences between groups

Two-way ANOVA, with ANOVA type III sum of squares analysis for unbalanced design (i.e., unequal number of participants in each group), was implemented to explore the differences in G-SAP-PD subscale scores between PwP+FOG and PwP-FOG. We found a significant main effect of freezing frequency (F[3]= 32.87, ƞ^2^=0.17, p<0.001) and a significant subscale X freezing frequency interaction (F[9]= 2.63, ƞ^2^=0.01, p=0.005), but no main effect of G-SAP-PD subscale (p=0.092). Post-hoc tests (Tukey) showed that across subscales, transformed G-SAP-PD scores were significantly higher for each subgroup of PwP+FOG compared to patients who never experience freezing, with highest scores for freezes everyday group (mean overall Z-scores: PwP-FOG -0.353±0.91, freezes hardly ever 0.24±0.90, freezes most weeks 0.42±0.92, and freezes everyday 0.80±0.93, p<0.001). Furthermore, we found no differences between subscale scores within each freezing frequency subgroup (p’s>0.23), except for significantly higher Rumination compared to conscious movement processing in the freezing everyday group (1.21±0.90 vs. 0.58±0.75, p=0.039). Results are summarized in Figure 2.

**Figure 2:**
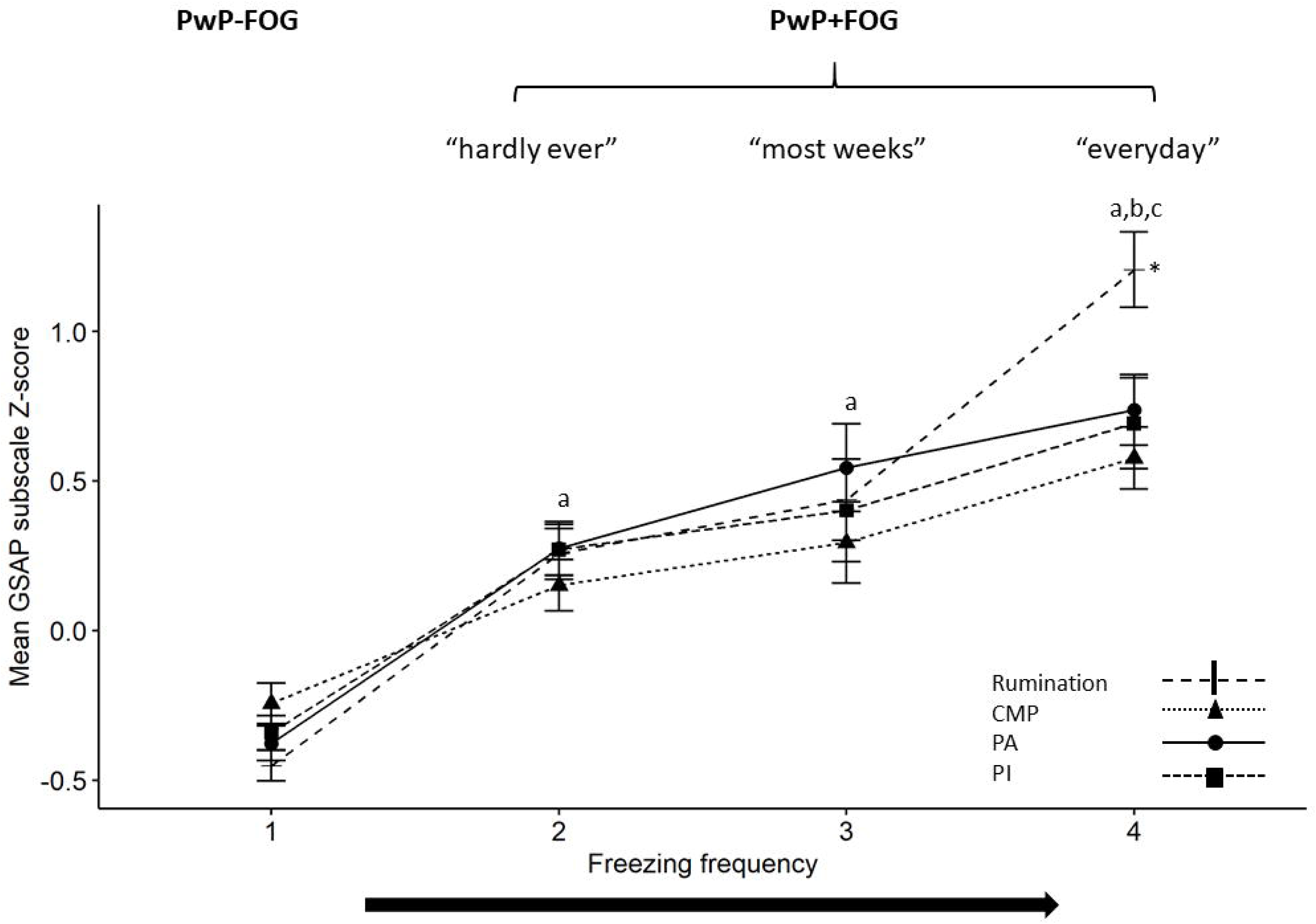
Mean (± standard error) Z-scores for each of the four subscales of the Gait-Specific Attentional Profile. Data are presented separately as a function of self-reported frequency of freezing of gait. Note: homogeneity of variances was assessed by plotting the residuals against the fitted values. Since there was no apparent relationship between the two, homogeneity was assumed. However, Levene’s test and Shapiro-Wilk test were significant (p=0.008 and p<0.001, respectively), hence not supporting homogeneity and normality assumptions (see Figure S1 in *Supplementary materials* for details). This should be considered when interpreting the results. a- significantly greater overall score than PwP-FOG group, p<0.001 b- significantly greater overall score than freezing ‘hardly ever’ group, p<0.001 c- significantly greater overall score than freezing ‘most weeks’ group, p<0.001 * Significantly different from CMP, p=0.039 CMP = Conscious movement processing, PA = Physiological Arousal, PI = Processing Inefficiency.

### G-SAP-PD scores - association with frequency of freezing

Ordinal regression results are presented in Table 2. Twenty-six participants with missing responses for age, years since diagnosis or experiencing balance problems, were not included in the regression analysis. Of the G-SAP-PD subscales, only higher Rumination subscale scores (OR=1.323, [1.214, 1.440]) and higher Physiological Arousal subscale scores (OR=1.195, [1.037, 1.377]) were associated with significantly greater odds of experiencing more frequent freezing. With regard to the control variables, only years since diagnosis was significantly associated with freezing frequency (OR=1.140, 95% CI = [1.091, 1.192]).

**Table 2.** Results of ordinal regression analysis of G-SAP-PD scores as a function of freezing of gait frequency.^a^.

| | OR [95% CI] | Wald $\chi^2$<br>(df=1) | *p* |
| --- | --- | --- | --- |
| Age in years | 0.988 [0.964, 1.012] | 0.942 | 0.332 |
| Years since diagnosis | 1.140 [1.091, 1.192] | 33.329 | **<.001** |
| Processing Inefficiency | 1.124 [0.991, 1.274] | 3.329 | 0.068 |
| Physiological Arousal | 1.195 [1.037, 1.377] | 6.005 | **0.014** |
| Rumination | 1.323 [1.214, 1.440] | 40.944 | **<.001** |
| Conscious Movement Processing | 0.961 [0.873, 1.056] | 0.684 | 0.408 |
| Balance/gait problems<sup>b</sup> | 0.559 [0.308, 1.014] | 3.669 | 0.055 |

NB: OR = odds ratio, values>1 indicate increase in odds of experiencing more frequently freezing; df=degrees of freedom; Model-parameters: Improvement in fit vs. intercept-only model ( $\chi^2=201.903$ , df=7, $p<0.001$ ); Goodness-of-fit indices: Pearson ( $\chi^2=1120.421$ , df=1217, $p=0.977$ ), Deviance ( $\chi^2=720.290$ , df=1217, $p=1.000$ ); Nagelkerke pseudo $R^2=0.435$ .

### G-SAP-PD scores cut off for predicting freezing

Since Rumination had the largest effect size for the difference between PwP+FOG and PwP-FOG and the highest odds ratio in the ordinal regression this sub-scale was used in the ROC analysis. Figure 3 presents the ROC curve for predicting FOG (Figure 3A) and freezing every day (Figure 3B). AUC was 0.777 and 0.854, for distinguish PwP-FOG from PwP+FOGand freezing every day, respectively, indicating good diagnostic accuracy. Rumination subscale cut-offs scores of 6.5 and 9.5 were found to be the optimal cut-offs (i.e., highest Youden index) for classifying PwP+FOG vs PwP-FOG and for freezing ‘every day’, respectively (Table S3 in Supplementary materials).

**Figure 3:**
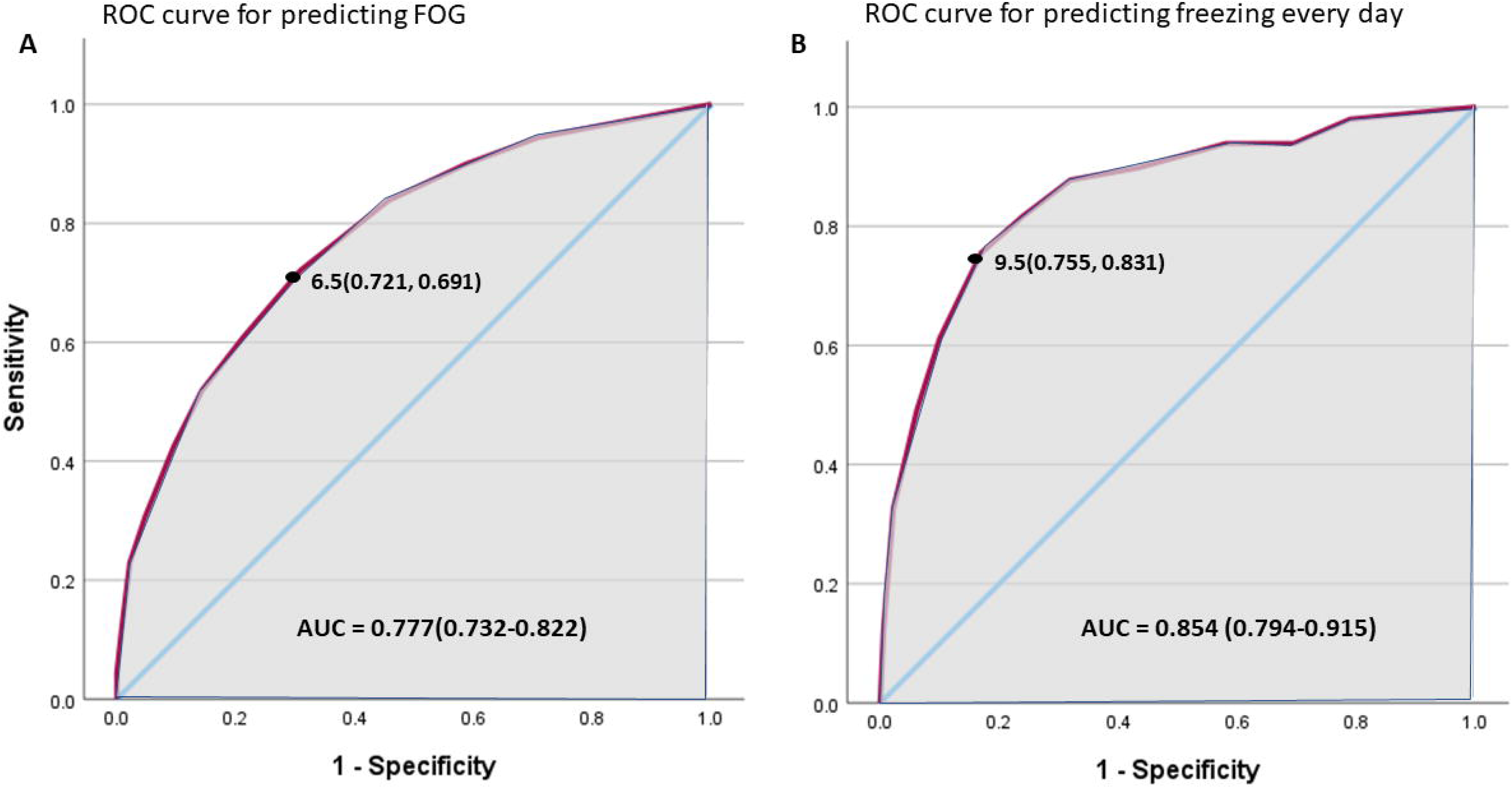
Receiver Operating Characteristic (ROC) curve (solid line) for predicting FOG (A) or freezing every day (B) using the G-SAP-PD, Rumination sub-scale. Diagonal line represents a random prediction. Grey area represents the area under curve (AUC) and black point represents the cut-off point with sensitivity and specificity in parenthesis.

## Discussion

This study supports the validity and internal consistency of the G-SAP-PD to measure attentional factors (Physiological Arousal, Rumination, CMP and Processing Inefficiencies) implicated in influencing FOG in PwP. Overall, confirmatory factor analysis and measurement invariance testing showed that the G-SAP-PD has a similar four factor structure as the original scale validated in community-dwelling older adults[18], with each of these subscales also showing sufficient internal consistency. Further, our results show that these G-SAP-PD scores may provide insight into how anxiety-related attentional characteristics may influence FOG frequency. Finally, ordinal regression revealed significant associations between FOG frequency and both Physiological Arousal and Rumination G-SAP-PD subscales, with strongest associations reported for the latter. Combined with findings of greater covariance between Physiological Arousal and CMP in PwP-FOG compared to PwP+FOG, and significantly higher Rumination scores compared to CMP scores among PwP with the most frequent freezing (i.e., on a daily basis), we therefore propose that Rumination contributes to freezing through disruption of conscious goal-directed behaviour that typically is required to maintain motor performance among PwP. We established cut-offs for scores on the Rumination subscale to distinguish PwP-FOG from PwP+FOG (cut-off: 6.5), and to distinguish those with most frequent freezing from those with less frequent freezing (cut-off: 9.5).

### G-SAP-PD reliability

The G-SAP scale was originally developed and validated in healthy older adults with the four attentional subscales selected due to their implication in influencing the control of balance and gait and therefore fall-risk[18]. Our results support the reliability of the G-SAP-PD to measure these same constructs in PwP (Physiological Arousal, Rumination, CMP and Processing Inefficiencies). It is important to note that researchers using the original G-SAP scale (https://osf.io/n7rcm/) to evaluate PwP, can adapt outcomes to the adapted version presented here (i.e., G-SAP-PD), by simply removing item A2 from the subscale of Physiological Arousal (formerly labelled as ‘Anxiety’).

### Associations between FOG severity and attention related cognitive processes

ACT describes how anxiety may shift cognitive processes away from <u>goal-directed</u> (dorsal) attentional system to information processing and associated <u>stimulus-driven</u> attentional system[10]. This phenomenon is apparent in PwP and highlighted in our findings.

#### Rumination

Researchers have previously demonstrated a relationship between physiological arousal and FOG[49], typically inferred through observations of altered heart rate and/or skin conductance around FOG onset[49,50]. This relationship is unlikely to be linear. Indeed, traditional conceptualisations of the relationship between arousal and motor performance (e.g., Yerkes-Dodson’s Law[51]) strongly implicate that an optimum level of arousal exists for a given task and that this might fluctuate based on several factors, such as individual characteristics of the performer. This notion has already been proposed in the specific context of FOG[52]. It infers that people may require a certain level of arousal to allocate attention in a manner that is sufficient to compensate for deficient automatic motor control processes. However, heightened arousal beyond this point is likely to compromise movement. Given existing literature[12–14,53] and conceptualisations in older adults with concerns about falling[18], we suggest that emergence of cognitive anxieties (i.e., rumination) could be particularly problematic. The primary reason for this is that worrisome thoughts, while potentially related to adaptive decision-making around a given task (e.g., opting to use a walking stick or not), are irrelevant to current motor output and therefore likely to add unnecessary cognitive demands that exacerbate attention related cognitive processing inefficiencies. The cross-talk model suggests these may culminate in increasing limbic load (secondary to higher levels of anxiety and competing inputs) which in turn could provoke FOG by overloading the striatum, interfering with normal basal ganglia motor processing[54]. As such, worries experienced in daily life could serve as distractions from consciously processing movements in much that same way as that observed in laboratory-based studies using cognitive dual-tasking paradigms that are known to exacerbate FOG[32,55]. However, change in *physiological* arousal/*somatic* anxiety, as outlined by the ACT[10], is unlikely to be the sole anxiety-related variable that exacerbates FOG in daily life. Our data suggest that a marker of *cognitive* anxiety (namely worrisome thoughts as measured using the Rumination subscale) is substantially increased in people with frequent FOG and is the factor most strongly associated with FOG severity (independently from Physiological Arousal).

#### Physiological Arousal and CMP

In healthy populations performing skilled motor tasks, such as in the context of sport, military or surgery, conscious control of movement can compromise performance by virtue of an over-reliance on explicit knowledge and reduction in implicit/automatic control processes[56]. However, the adaptive role of CMP has been demonstrated in people whose automaticity is compromised to some extent, such as older adults with reduced functional balance or stroke survivors[57,58]. We argue that these observations might be extrapolated to the context of Parkinson’s, where basal ganglia impairments cause pronounced deficiencies in automaticity, thereby increasing reliance on CMP as a necessary compensatory mechanism. Indeed, in the current study, we found evidence to suggest that PwP-FOG may have been better able to offset negative effects of arousal compared to PwP+FOG by engaging in increased CMP, as the PwP-FOG demonstrated greater covariance between Physiological Arousal and CMP. It might be the case that PwP+FOG are already highly engaged in CMP regardless of their anxiety level, potentially reflecting more progressed disease, and hence, greater deficiencies in automaticity that need to be compensated. This could potentially create a ceiling effect, where there is limited scope for investing greater CMP. This might be an explanation for the lower CMP and arousal covariance for the PwP+FOG.

### Anxiety management in the context of FOG

The understanding of the relationship between anxiety and FOG in recent years along with growing awareness of anxiety’s detrimental effect on quality of life[59,60] has led to suggestions that dysfunction in arousal and anxiety-related attentional processes could provide a useful biomarker for early identification of PwP who might develop FOG[8,33]. Apart from targeting these cognitive processes themselves, anxiety management of PwP may therefore be a fruitful avenue for alleviating some of the motor symptoms associated with Parkinson’s. However, effective strategies for managing anxiety and its influence on motor symptoms are currently insufficient[61]. Typically, treatment approaches are pharmacological, involving drugs like selective serotonin reuptake inhibitors, benzodiazepines and antidepressant drugs[59,62]; and non-pharmacological, including cognitive behavioural therapy, transcranial magnetic stimulation, yoga and breathing control, to name a few[62,63]. While a range of interventions have been presented, current evidence reporting the effectiveness of existing treatment options is weak[59,64–66]. In a recent paper, Hinkle et al.[67], found a weak association between improved anxiety scores and change in motor symptoms in response to dopamine intake. Further study is warranted to decipher the neurophysiological and neurobiological associations between dysfunction in brain networks associated with arousal and cognition. However, since we don’t fully understand how anxiety effects PwP and its relation to FOG it is essential that the lived experiences of anxieties relating to FOG are considered within these endeavours and used in the design of non-pharmacological treatments. Ultimately, it would be highly valuable if this could lead to interventions and/or resources that help PwP to self-manage anxiety.

## Limitations

The current study has several limitations. First, all measures were self-reported and we therefore could not independently verify participants’ FOG status. However, the current approach did allow us to recruit a very large sample of PwP from across the UK, which is essential to perform a comprehensive scale validation . Nonetheless, it is possible that participants may have misunderstood certain questions or provided biased answers, given previous evidence of inaccuracies in self-reported levels of physical activity[68] and FOG frequency[69,70]. That being said, the pattern of results presented here fits with previous reports (e.g., associations between freezing status and frequency with balance problems, years since diagnosis, etc), and there are no clear additional sources of bias within our protocol that might compromise self-reported outcomes presented here[71].

As our study is cross-sectional in nature, inferences regarding potential *causal* links between G-SAP-PD subscales and FOG cannot be made, and require further study.

## Conclusion

Th G-SAP-PD questionnaire could be used to monitor attentional constructs related to FOG in PwP. Our data suggests that perceptions of physiological arousal are associated with potentially more adaptive CMP in PwP-FOG. Conversely, PwP+FOG appear to demonstrate anxiety-related vulnerabilities characterised by a relative inability to engage in compensatory goal-directed focus of attention, potentially driven by heightened stimulus-directed attention and associated worrisome thoughts. It is important to consider that the relationships between physiological arousal, CMP, and rumination were observed when controlling for disease duration and self-reported balance impairment. While previous reports of higher G-SAP sub-scale scores in PwP+FOG were interpreted as potentially being driven by longer disease duration[37], the current data indicate that the relationship between physiological arousal and ruminations in PwP with frequent freezing could be driven, at least in part, by worrisome thoughts exacerbating FOG. Future research should investigate the interplay between worrisome thoughts and FOG. After all, tendencies in the way people allocate attention prior to or during FOG are perhaps more readily modifiable compared to progressive and chronic changes in automaticity/attentional capacity. Further study is required to test these ideas.

## Acknowledgements

The authors will like to acknowledge the participants in the study that contributed from their valuable time for the successes of our work.

## Declarations

### Availability of data and material

The datasets during and/or analysed during the current study available from the corresponding author on reasonable request.

### Consent for publication

“Not applicable”

### Authors’ contributions

UR analysed and interpreted the patient data, and drafted the manuscript. AC contributed to the inception of the study and collected the data. MN contributed to the inception of the study and was a contributor in writing the manuscript. EK was a major contributor to the data analysis and interpretation and in writing the manuscript. WY conceptualized and advised on the study as the main PI, was a major contributor in collecting the data and writing the manuscript. All authors read and approved the final manuscript.

### Supplementary Material

Supplementary materials include the G-SAP-PD form and elaborations on the result section.

## List of abbreviations

(G-SAP-PD): Adapted Gait Specific Attentional Profile
(G-SAP): Gait-Specific Attentional Profile
(FOG): freezing of gait
(PwP): people with Parkinson’s
(CMP): Conscious Movement Processing
(PwP+FOG): People with Parkinson’s with FOG
(PwP-FOG): People with Parkinson’s without FOG
(ACT): Attention Control Theory
(PI): processing inefficiencies
(CFA): confirmatory factor analysis
(GFI): goodness-of-fit index
(CFI): comparative fit index
(SRMR): standardized root mean squared residual
(RMSEA): root mean square error of approximation
(AUC): area under the curve
(OR): Odds Ratio
(ROC): Receiver Operating Characteristic

