## Supplementary Materials for "Anxiety-related attentional characteristics and their relation to freezing of gait in people with Parkinson’s – cross-validation of the Adapted Gait Specific Attentional Profile (G-SAP-PD)"

**Methods:**

### Gait-Specific Attentional Profile-PD©

Date……………………………

**Mark the appropriate circle to indicate**

**<u>how you feel when you walk</u>**

| | NOT AT ALL | NOT VERY MUCH | MODERATELY SO | OFTEN | VERY MUCH SO |
|---|---|---|---|---|---|
| A1. I feel strained……………………………………………………………………………… | ① | ② | ③ | ④ | ⑤ |
| A3. I think about previous occasions when I lost my balance…………. | ① | ② | ③ | ④ | ⑤ |
| A4. I think about what would happen if I fell………………………………….. | ① | ② | ③ | ④ | ⑤ |
| A5. I get confused and make illogical decisions……………………………. | ① | ② | ③ | ④ | ⑤ |
| A6. Worrisome thoughts about falling run through my mind………… | ① | ② | ③ | ④ | ⑤ |
| A7. I try to think about the way I walk/move…………………………………. | ① | ② | ③ | ④ | ⑤ |
| A8. I consciously try to control my movements………………………………. | ① | ② | ③ | ④ | ⑤ |
| A9. I examine the way I walk/move………………………………………………… | ① | ② | ③ | ④ | ⑤ |
| A10. I feel tense……………………………………………………………………………… | ① | ② | ③ | ④ | ⑤ |
| A11. I find it difficult to concentrate on two things at once……………. | ① | ② | ③ | ④ | ⑤ |

**To be completed by the researcher/clinician**

**Calculate the total score from each item relating to the four categories:**

Physiological arousal (sum of A1, A10) = ………………

Conscious movement processing (sum of A7, A8, A9) = ………………

Task-irrelevant ruminations/thoughts (sum of A3, A4, A6) = ………………

Processing inefficiencies (sum of A5, A11) = ………………

**Results:**

**Table S1.** Results of measurement invariance testing.

| Invariance test | $\chi^2$ | CFI<br>GFI | RMSEA (90%CI) | SRMR | Model comp. | $\Delta\chi^2$ | $\Delta$CFI<br>$\Delta$GFI | $\Delta$RMSEA<br>$\Delta$SRMR | Decision |
|---|---|---|---|---|---|---|---|---|---|
| **1. Config.** | 123.330<br>df=58<br>*p*<0.001 | 0.965<br>0.946 | 0.051<br>[0.039,<br>0.064] | 0.043 | N/A | N/A | N/A | N/A | Accept |
| **2. Metric** | 126.262<br>df=64<br>*p*<0.001 | 0.967<br>0.945 | 0.048<br>[0.035,<br>0.060] | 0.044 | 1 | 2.932<br>df=6<br>*p*=0.817 | 0.002<br>-0.001 | -0.003<br>0.001 | Accept |
| **3. Scalar** | 160.288<br>df=76<br>*p*<0.001 | 0.954<br>0.931 | 0.052<br>[0.041,<br>0.063] | 0.060 | 2 | **34.025**<br>**df=10**<br>***p*<0.001** | -0.013<br>-0.014 | 0.004<br>**0.016** | (Accept)* |
| **3a. Partial Scalar**** | 149.564<br>df=73<br>p<0.001 | 0.959<br>0.935 | 0.050<br>[0.038,<br>0.061] | 0.054 | 3 | **23.302**<br>**df=9**<br>***p*=0.006** | -0.008<br>-0.010 | 0.002<br>0.011 | (Accept)* |
| **3b. Partial Scalar***** | 142.852<br>df=72<br>p<0.001 | 0.962<br>0.937 | 0.048<br>[0.036,<br>0.059] | 0.055 | 3a | **16.590**<br>**df=8**<br>***p*=0.035** | -0.005<br>-0.008 | 0.000<br>0.012 | (Accept)* |

**NB:** CFI = Comparative fit index; Config. = Configural; GFI = Goodness-of-fit index; Model comp. = Model comparison; N/A= Not applicable; RMSEA = Root mean square error of approximation; SRMR = Standardized root mean squared residual; df = degrees of freedom; Model fit indices that exceed the threshold for acceptable model fit change are emphasized;

* Scalar invariance was partly confirmed: $\Delta$CFI, $\Delta$GFI, $\Delta$RMSEA were acceptable, but $\Delta\chi^2$ was significant and $\Delta$SRMR>0.015;

** Backward releasing of constraints revealed that allowing the covariance for ‘Physiological Arousal’ and ‘Conscious Movement Processing’ (model 3a) to differ across groups resulted in improved fit across all indices, except that $\Delta\chi^2$ remained significant;

*** Additional releasing of constraints related to the variance of scores on the Conscious Movement Processing subscale (model 3b) resulted in further significantly improved fit across all indices. While $\Delta\chi^2$ remained significant, releasing of other constraints did not significantly improve model fit further.

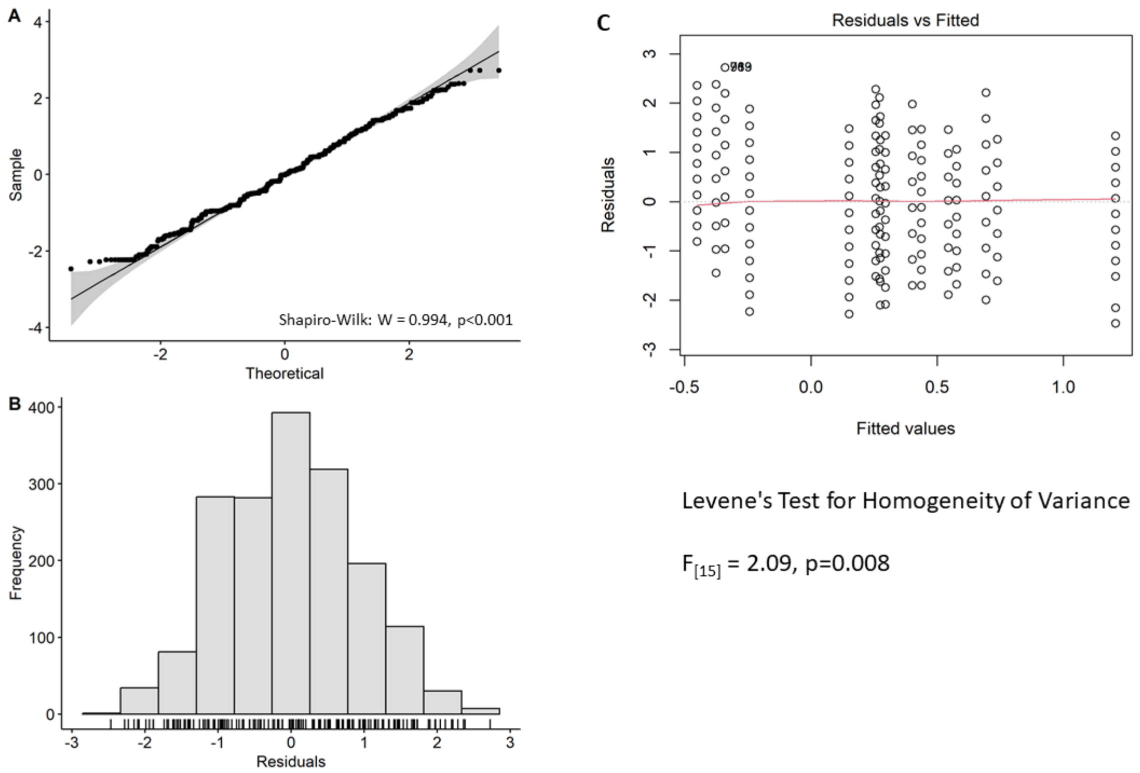


**Figure S1:** Summery of assumptions testing for 2-way ANOVA where (A) is a Q-Q plot of ANOVA residuals (black dots) in relation to the identity line (y = x, black solid line) and their 95% confidence interval in shaded grey; (B) histogram of the residuals; and (C) Scatter plot of the residuals against the fitted values with a regression line (red solid line) for assessing homogeneity of variances.

***G-SAP-PD scores - association with frequency of freezing***

Multinomial logit model is summarized in Table S2. The analysis revealed that all levels of frequency of freezing were significantly associated with years since diagnosis (Wald $\chi2$ = 5.393-26.217, p's≤0.020) and Rumination (Wald $\chi2$ = 6.859-35.535, p's≤0.009), such that the odds for a one-unit increase in the variable years since diagnosis is 1.084 for being in Hardly ever group vs. the Never group, 1.208 for being in Most weeks group vs. the Never, and 1.238 for being in the  Everyday group vs. the Never group. The relative risk for a one-unit increase in the variable rumination is 1.218 for being in Hardly ever group vs. the Never group, 1.241 for being in Most weeks group vs. the Never, and 1.797 for being in the Everyday group vs. the Never group.

The Hardly ever group also showed significant associations with Age in years and Processing Inefficiency (Wald $\chi2$ = 6.948, p=0.008, Wald $\chi2$ = 4.267, p=0.039, respectively), such that the odds of being a year older is 0.959 and the relative risk for scoring one point higher on the Processing Inefficiency subscale is 1.192 for being in the Hardly ever group vs. the Never group. The Most weeks group also showed significant association with Anxiety (Wald $\chi2$ = 9.458, p=0.002) such that the relative risk for a one-unit increase in the variable anxiety is 1.543 for being in Most weeks group vs.

the Never group. The Everyday group also showed significant association with Balance/gait problems (Wald $\chi2$ = 3.893, p=0.049) such that the relative risk ratio for developing a Balance/gait problem is 0.102 for being in the Everyday group vs. the Never group.

**Table S2.** Results of multinomial logit model regression analysis of G-SAP-PD scores and freezing of gait frequency**.**

| **Freezing Frequency**[a] | | **OR [95% CI]** | **Wald $\chi^2$ (df=1)** | ***p*** |
|---|---|---|---|---|
| **Everyday** | Age in years | 1.042[0.990, 1.097] | 2.434 | .119 |
| | Years since diagnosis | 1.238[1.141, 1.343] | 26.217 | **<.001** |
| | Processing Inefficiency | 1.177[0.931, 1.486] | 1.859 | .173 |
| | Anxiety | 1.208[0.919, 1.587] | 1.841 | .175 |
| | Rumination | 1.797[1.482, 2.178] | 35.535 | **<.001** |
| | Conscious Movement Processing | 1.016[0.825, 1.250] | 0.021 | .884 |
| | Balance/gait problems[b] | 0.102[0.011, 0.985] | 3.893 | **.049** |
| **Most weeks** | Age in years | 0.963[0.918, 1.010] | 2.460 | .117 |
| | Years since diagnosis | 1.208[1.112, 1.313] | 19.790 | **<.001** |
| | Processing Inefficiency | 1.196[0.945, 1.514] | 2.211 | .137 |
| | Anxiety | 1.543[1.170, 2.035] | 9.458 | **.002** |
| | Rumination | 1.241[1.056, 1.459] | 6.859 | **.009** |
| | Conscious Movement Processing | 0.907[0.748, 1.099] | 0.995 | .319 |
| | Balance/gait problems[b] | 0.965[0.315, 2.963] | 0.004 | .951 |
| **Hardly ever** | Age in years | 0.959[0.929, 0.989] | 6.948 | **.008** |
| | Years since diagnosis | 1.084[1.013, 1.161] | 5.393 | **.020** |
| | Processing Inefficiency | 1.192[1.009, 1.409] | 4.267 | **.039** |
| | Anxiety | 1.162[0.966, 1.399] | 2.529 | .112 |
| | Rumination | 1.218[1.091, 1.360] | 12.310 | **<.001** |
| | Conscious Movement Processing | 0.917[0.815, 1.030] | 2.129 | .145 |
| | Balance/gait problems[b] | 0.543[0.273, 1.082] | 3.011 | .083 |

NB: OR = odds ratio, values>1 indicate increase in odds of experiencing more frequently freezing; df=degrees of freedom; Model-parameters: Improvement in fit vs. intercept-only model ($\chi^2$=235.335, df=21, $p$<0.001); Goodness-of-fit indices: Pearson ($\chi^2$=1193.434, df=1203, $p$=0.572), Deviance ($\chi^2$=686.858, df=1203, $p$=1.000); Nagelkerke pseudo $R^2$=0.489.

a. The reference category is: Never.
b. Reference category is the self-reported problems with balance or gait group (N=293).

***G-SAP-PD scores cut off for predicting freezing***

**Table S3:** Measures of Diagnostic Accuracy from the Receiver Operating Characteristic (AUC) Curve Predicting FOG and freezing every day from G-SAP-PD, Rumination sub-scale.

| | cut-off score | Sensitivity | Specificity | Youden's Index | AUC (95%CI)<br>P<br>Gini Index |
|---|---|---|---|---|---|
| **PwP+FOG** | | | | | 0.777 (0.732-0.822)<br><0.001<br>0.554 |
| Optional cut-off 1 | 5.5 | 0.838 | 0.543 | 0.381 | |
| Optional cut-off 2 | **6.5** | **0.721** | **0.691** | **0.412** | |
| Optional cut-off 3 | 7.5 | 0.609 | 0.787 | 0.396 | |
| **PwP+FOG everyday** | | | | | 0.854 (0.794-0.915)<br><0.001<br>0.709 |
| Optional cut-off 1 | 8.5 | .816 | 0.761 | .577 | |
| Optional cut-off 2 | **9.5** | **.755** | **0.831** | **.586** | |
| Optional cut-off 3 | 10.5 | .612 | 0.900 | .512 | |

Abbreviations: PwP+FOG = People with Parkinson's that experience freezing of gait.
